# Rapid evolution of female-biased genes among four species of *Anopheles* malaria mosquitoes

**DOI:** 10.1101/081620

**Authors:** Francesco Papa, Nikolai Windbichler, Robert M. Waterhouse, Alessia Cagnetti, Rocco D’Amato, Tania Presampieri, Mara K. N. Lawniczak, Tony Nolan, Philippos Aris Papathanos

## Abstract

Understanding how phenotypic differences between males and females arise from the sex-biased expression of nearly identical genomes can often reveal important insights into the biology and evolution of a species. Among *Anopheles* mosquito species, these phenotypic differences include vectorial capacity, as it is only females that blood feed and thus transmit human malaria. Here, we use RNA-seq data from multiple tissues of four vectors spanning the *Anopheles* phylogeny to explore the genomic and evolutionary properties of sex-biased genes. We find that in these mosquitoes, in contrast to what has been found in many other organisms, female-biased genes are more rapidly evolving in sequence, expression, and genic turnover, than male-biased genes. Our results suggests that this atypical pattern may be due to the combination of sex-specific life history challenges encountered by females, such as blood feeding. Furthermore, female propensity to only mate once in nature in male swarms likely diminishes sexual selection of post-reproductive traits related to sperm competition among males. We also develop a comparative framework to systematically explore tissue- and sex-specific splicing, to document its conservation throughout the genus and identify a set of candidate genes for future functional analyses of sex-specific isoform usage. Finally, our data reveals that the deficit of male-biased genes on the X chromosomes in *Anopheles* is a conserved feature in this genus and can be directly attributed to chromosome-wide transcriptional regulation that demasculinizes the X in male reproductive tissues.

## Introduction

One of the mot extreme examples of phenotypic variation between individuals of the same species is the difference between the sexes. However, apart from sex-limited Y and W chromosomes that are typically gene poor and small, males and females share a common genome. Sexual dimorphism is therefore largely achieved through sex-biased expression or splicing of this shared genome throughout development (Connallon and Knowles 2005; Ellegren and Parsch 2007). Genome sharing however, imposes evolutionary constraints on chromosomal and genic content, because these must respond to different regimes of natural and sexual selection in each sex. Mutations that lead to changes in gene expression or coding sequence can even result in conflicting fitness optima - beneficial for one sex but detrimental to the other - a process called sexual antagonism (Rice 1996, 1998; Chippindale et al. 2001; Chapman et al. 2003).

A common pattern that has emerged from comparative genomic studies of sex-biased gene expression in insects (Zhang et al. 2004, 2007; Assis et al. 2012; Wang et al. 2015), worms (Reinke et al. 2004; Cutter and Ward 2005), birds (Harrison et al. 2015), rodents (Torgerson et al. 2002; Good and Nachman 2005) and primates (Khaitovich et al. 2005), has been that male-biased genes more common, more tissue-specific and more abundantly expressed than female-biased genes (Grath and Parsch 2016). Analyses of their evolutionary dynamics have also shown that male-biased genes are typically more rapidly evolving than female-biased genes: they display higher birth and death rates (turnover), higher rates of expression divergence and sex-bias switching and a lower conservation of protein sequences. In particular, genes expressed during spermatogenesis or in tissues that produce the seminal fluids transferred to females during mating are extremely evolutionarily dynamic, emphasizing that male-male and sperm competition are strong forces driving genome evolution (Ellegren and Parsch 2007).

To date, sex-biased gene expression and its evolutionary consequences have mainly been explored in species in which the majority of phenotypic variation between the sexes is centered on the act of mating, rather than substantial divergence in other life history traits. In this regard, mosquito species may provide interesting insights for the study of sexual dimorphism within a more diverse context, as adult mosquitoes exhibit a rich repertoire of unique sex-specific biology related to host-seeking, blood-meal ingestion and encounters with parasites and pathogens. Females have a highly specialized apparatus for host seeking that responds to multiple sensory inputs including CO2, heat and olfactory cues (Sutcliffe 1994; van Breugel et al. 2015; Maekawa et al. 2011). In the case of a handful of the *Anopheles* species such as *An. gambiae* and *An. arabiensis*, their adaptation to preferentially seeking and biting human hosts underpins their role as vectors of malaria parasites and their enormous impact on public health (Garrett-Jones et al. 1980). Blood-feeding also means that female mouthparts differ structurally from those of males, which feed exclusively on nectar from flowers (Robinson 1939). In addition, the salivary glands produce a number of female-specific enzymes that are injected into the host epithelium to inhibit blood coagulation and digestion of the blood meal involves the expression of a diverse catalog of proteins and enzymes (Kalume et al. 2005; Fontaine et al. 2012; Arcà et al. 2005; Arcá et al. 1999). The digestive and immune systems of females are also focused on dealing with the specific challenges that arise from taking large blood meals that cause substantial changes in the microbiota community, both in terms of their diversity and abundance (Boissière et al. 2012; Wang et al. 2011; Pumpuni et al. 1996). Finally, males also engage in sex-specific behaviors, such as congregating in large swarms that act as their principal mating arenas (Butail et al. 2013).

How does this rich repertoire of sexual dimorphism in mosquitoes affect the evolutionary dynamics and abundance of sex-biased genes, especially in the light of the many differences that manifest in tissues of the carcass? Here we perform an analysis of the evolutionary histories and the signatures of selection on the genes expressed in a sex-biased manner to begin to address these questions. In previous work we have catalogued sex biased gene expression of *An. gambiae* in both adult and immature stages (Magnusson et al. 2011) and in dissected tissues using microarray platforms (Baker et al. 2011). However the extent and conservation of sex-biased gene expression in other *Anopheles* species is unknown, thus limiting the comparative analysis to species outside of the *Anopheles* genus. The recent completion of the genome sequencing of 16 additional *Anopheles* species (Neafsey et al. 2014) provides an opportunity to study the evolutionary consequences of sex-biased gene expression in much greater detail. Here we leverage these newly assembled resources and use transcriptome data based on Illumina RNAseq from four mosquito species spanning the genus, *An. gambiae*, *An. arabiensis*, *An. minimus* and *An. albimanus*. We selected these species because they span the entire phylogeny and because all are principal malaria vectors in the territories where they are endemic - Africa, Asia and Central America. We use these data to explore genome-wide evolutionary rates of sex-biased genes in carcass and reproductive tissues separately, and to determine whether their abundance and evolutionary properties reflect phenotypic and functional differences between the sexes.

## Results

### Transcriptome sequencing

Total mRNA was obtained from three biological replicates of dissected reproductive or carcass tissues of male and female adult pools from four malaria vectors that span the phylogeny of the *Anopheles* genus (Figure 1A and 1B). Reproductive tissues (RT) from females included the ovaries and the lower reproductive tract and for males included the testis and the accessory glands (Figure 1C). The remaining carcass (CA) included the head, thorax and abdomen. Females were blood-fed 48 hours prior to sampling to release ovarian development from its pre-vitellogenic diapause (Clements 1992). In total we generated 92.9 Gb of sequence read data in 100bp paired-end reads. After quality filtering, reads were mapped to the correct species assembled genomes retrieved from Vectorbase (Giraldo-Calderón et al. 2015) using Tophat2 (Kim et al. 2013) to quantify gene level expression. Expression values in read counts for each gene were extracted using HTSeq (Anders et al. 2014) and differential expression (DE) analysis was performed using DESeq-2 (Anders and Huber 2010) (Table S1). After DE analysis, the value of fragments per kilobase of transcript per million mapped reads (FPKM) for each gene were calculated and used for subsequent analysis.

**Figure 1.**
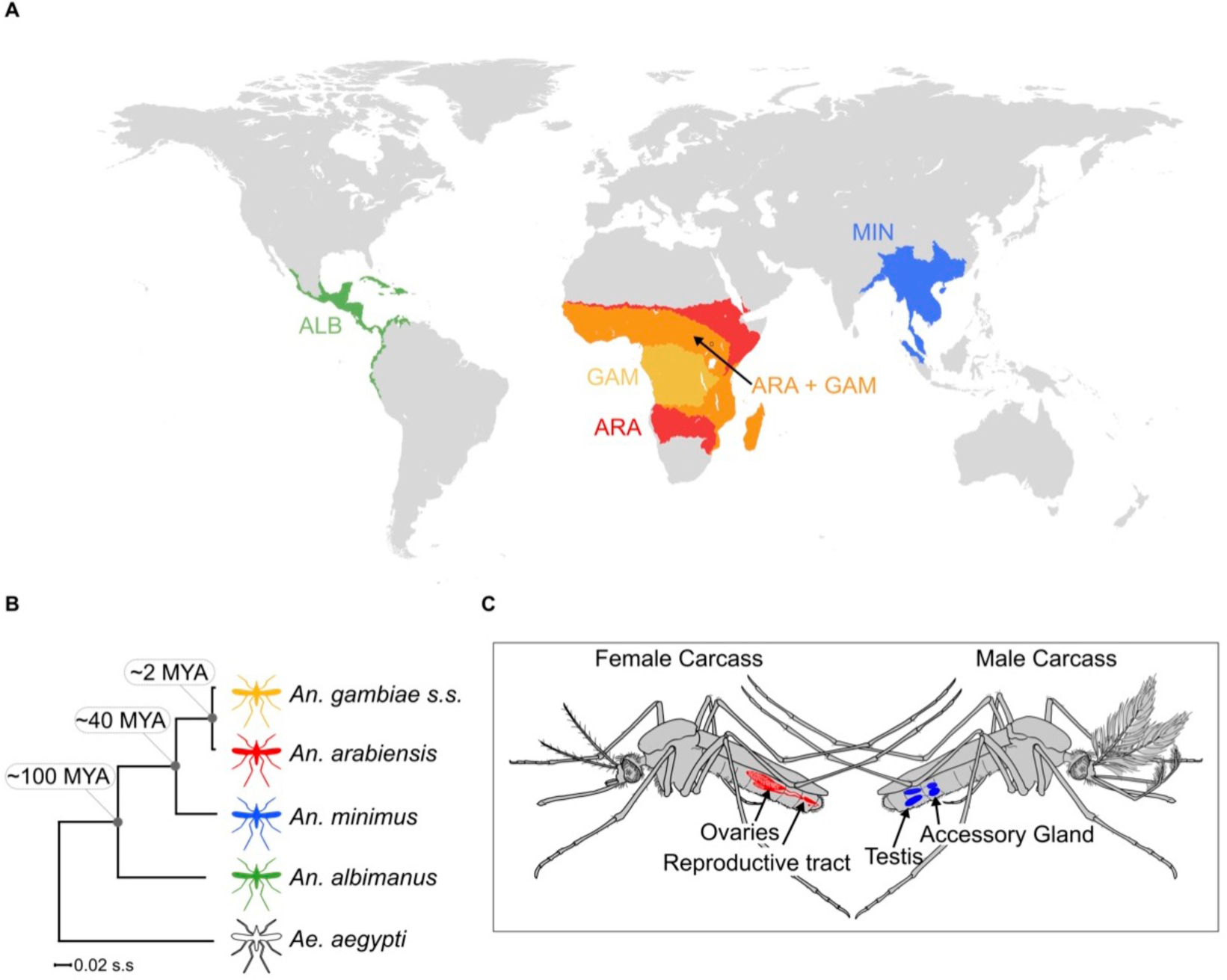
Study design: **A)** Global map displaying geographic ranges of the study species: *An. gambiae* (yellow), *An. arabiensis* (red) *An. minimus (*blue) *An. albimanus (*green). **B)** The maximum likelihood molecular phylogeneny of the study species within the *Anopheles* genus with *Aedes aegypti* as the outgroup. (Divergence time estimates from (Neafsey et al. 2014) and (Fontaine et al. 2015)) **C)** Overview of the dissected tissues used for RNAseq samples. Reproductive tissues included 48hrs post-blood meal ovaries and lower reproductive tract for females (in red) and testes with accessory glands for males (in blue). The remaining carcass tissues of both sexes are colored in grey.

### Cataloguing sex-biased gene expression

DE analysis was performed for each species in two steps: first testing for a sex-bias in gene expression between tissues (e.g. male CA versus female CA) and then a tissue-bias within each sex (e.g. male CA versus male RT) using a false discovery rate < 0.05 between conditions (Sup Figure1). Sex-biased genes that displayed significant (Bonferroni corrected p-value < 0.05) and concordant tissue-biases were classified into categories reflecting both traits (e.g female-biased between the RT samples and RT-bias between female tissues = female-RT-biased). Genes that displayed a sex-bias in both tissues and no bias between tissues in that sex were classified as ubiquitously sex-biased. Sex-biased genes that displayed conflicting sex- and tissue-bias (e.g. male-bias between RT samples but CA-bias within males) were not included in the final sex-biased categories. As a result we were able to specify for each differentially expressed gene a single sex-bias and tissue-bias call. On average 4290 genes per species (~33%) were classified as sex-biased in either one or both tissues. Sex-biased genes were then divided into one of three categories based on the magnitude of sex-biased expression: sex-specific, strong sex-bias (with a fold change (FC) greater than 5) and weak sex-bias (2>FC<5). To be classified as sex-specific, a gene had to display a significant sex-bias and a FPKM value lower than 10 in one sex but higher than 20FPKM in the other (see Methods for further details).

Consistent with findings in other species (Zhang et al. 2004, 2007; Reinke et al. 2004; Cutter and Ward 2005; Torgerson et al. 2002; Good and Nachman 2005; Khaitovich et al. 2005), we found that the majority of the sex-biased gene repertoire is derived from expression in the reproductive tissues, but in the four mosquito species analyzed, female-biased genes predominated over male-biased genes (Figure 2A). There was a higher correlation of levels of gene expression between the carcasses of the sexes (on average 0.7 Pearson correlation, p-value<2.2e-16) than between the reproductive tissues of the sexes (average 0.12 Pearson correlation and p-value<2.2e-16) (Figure 2B). Similar to other species, genes biased towards the male reproductive tissues were more likely to display tissue and sex specific expression compared to genes biased in the other three tissues (shown for *An. gambiae* in Figure 2C and remaining species in Sup. Figure2) and showed a higher magnitude of sex-bias (Figure 2E). To validate our DE classifications, we retrieved expression data for *An. gambiae* from the mosquito atlas (Mozaltas) study (Baker et al. 2011) - similar data are not available for the remaining species. We found that the majority of genes classified as sex-biased in the reproductive tissues were most abundantly expressed in the testis (50.4%), accessory glands (9.9%) and ovaries (30%), while carcass derived sex-biased genes were mainly composed of genes expressed abundantly in somatic tissues (Figure 2D). Our DE analysis and subsequent classification of sex and tissue biases were therefore consistent with results from previous experiments.

**Figure 2.**
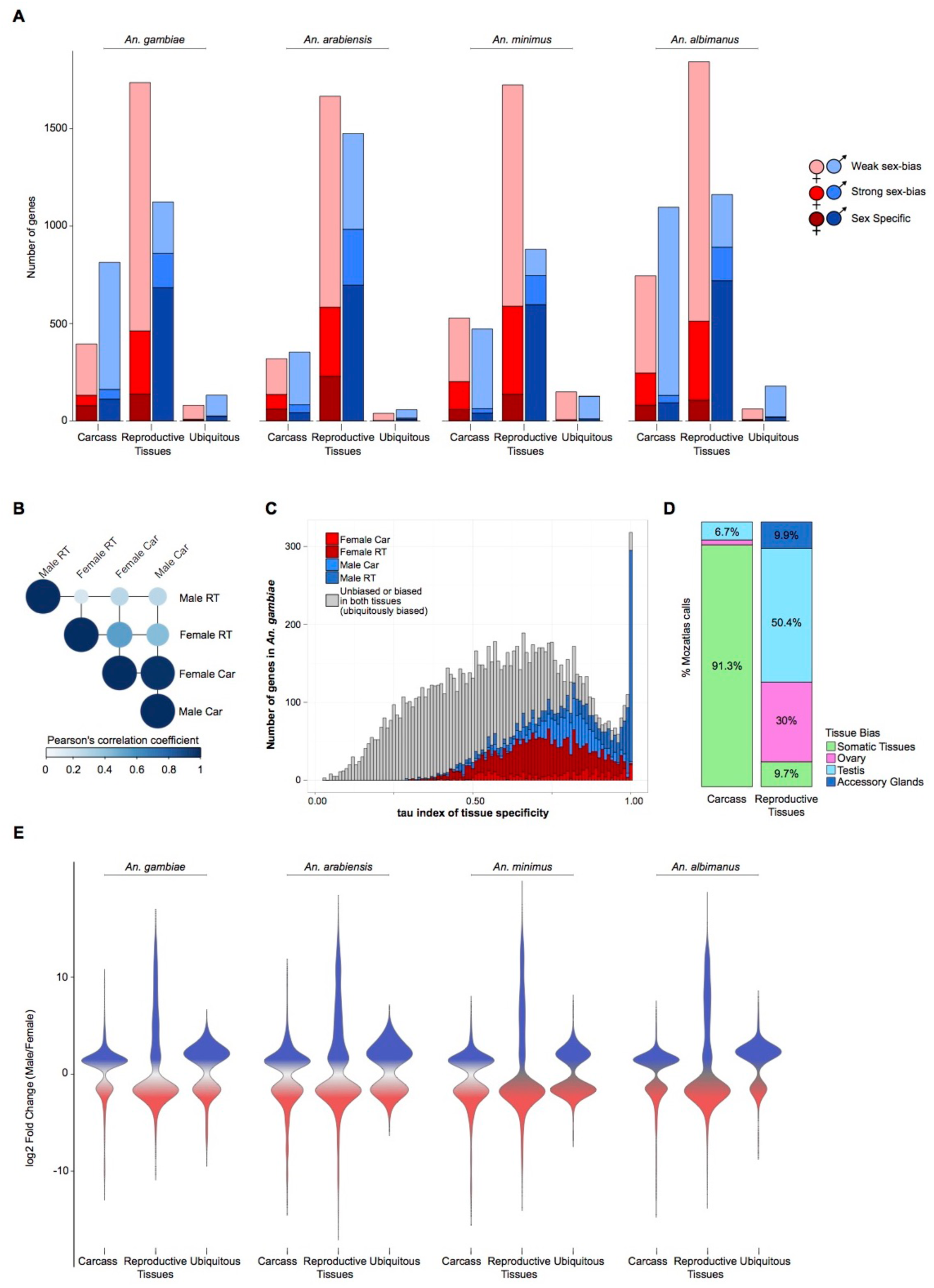
Sex-biased gene expression in the *Anopheles* genus. **A)** Total number of sex-biased genes in the four study species for each of three tissue expression biases: carcass, reproductive tissues and ubiquitously sex-biased. Female (red hues) and male (blue hues) sex-bias magnitudes are depicted as: dark - sex specific; medium - strongly sex-biased; light - weakly sex-biased. **B)** Average Pearson’s correlation coefficient of gene expression between sexed tissues in *An. gambiae*. **C)** Distribution of tau (*τ*) index of tissue specificity in *An. gambiae* for each sex- and tissue-biased expression classification, where increasing *τ* values respond to increased sample-specific expression. Genes with very high *τ* values (~1) are highly enriched for male reproductive tissue-biased genes. **D)** Comparison of sex-bias gene expression categories produced by this differential expression pipeline using RNA-seq with results from a previous microarray study for *An. gambaie* (MozAtlas (Baker et al. 2011)), showing strong congruence of gene expression bias in tissue classifications. **E)** Magnitude of gene expression ratios (log2 fold male/female) for each species. Male-biased genes of the reproductive tissues display the largest magnitude and range of biased expression.

### Evolution of sex biased gene expression

To study the molecular evolution of sex-biased gene expression in the *Anopheles* genus we compiled orthology relationships of all annotated genes of the four species and calculated expression and sequence divergence, as well as gene turnover rates separately for the carcass and the reproductive tissues. We catalogued 7400 one-to-one orthologs across the four species, referred to hereafter as the four-species orthologs (Table S2). Most of the analyses presented below are focused on analyses of these four-species orthologs, as this simplifies the interpretation of the results and because these are less likely to contain annotation errors that may bias data interpretation, for example incorrect automated gene predictions or orthology calls among many-to-many orthologs. However, because genes that are unambiguously orthologous across the entire genus may also show the least evolutionary divergence we also used a larger subset of orthologs with 31472 orthologs across all species, containing species-restricted genes (2289 on average per species) and those with one-to-one orthology between at least two of the study species (3183 orthogroups). Many-to-many orthologous relationships were excluded from the analysis.

#### Expression divergence

To study how sex-biased gene expression correlated with expression divergence we calculated the median standard deviation (SD) of the sex expression ratio (log2 male/female) between the four-species orthologs for each sex-bias classification (including unbiased), similarly to other studies (Zhang et al. 2007). The SD was calculated between orthologs of genes displaying significant sex-biased expression in each species and tissue, and values of expression ratios were drawn either from carcass or RT data depending on the sex- and tissue-bias classification of each gene. Overall, female-specific genes expressed in the carcass exhibited the highest levels of expression divergence of all sex-bias categories in all species (Figure 3A). In the carcass, expression divergence levels among sex-biased genes were higher than unbiased genes regardless of the direction and magnitude of the sex-bias (Wilcoxon rank-sum test p< 0.001) (Figure 3A). In the reproductive tissues, sex-biased genes exhibited greater dynamic ranges of SD compared to genes biased in the carcass (Figure 3B) and female-specific, male-specific and strongly male-biased genes showed high levels of expression divergence while all other classes evolve at similar or lower levels compared to unbiased genes (Figure 3A). These patterns were consistent in each of the four species individually (Sup. Figure 3A).

**Figure 3.**
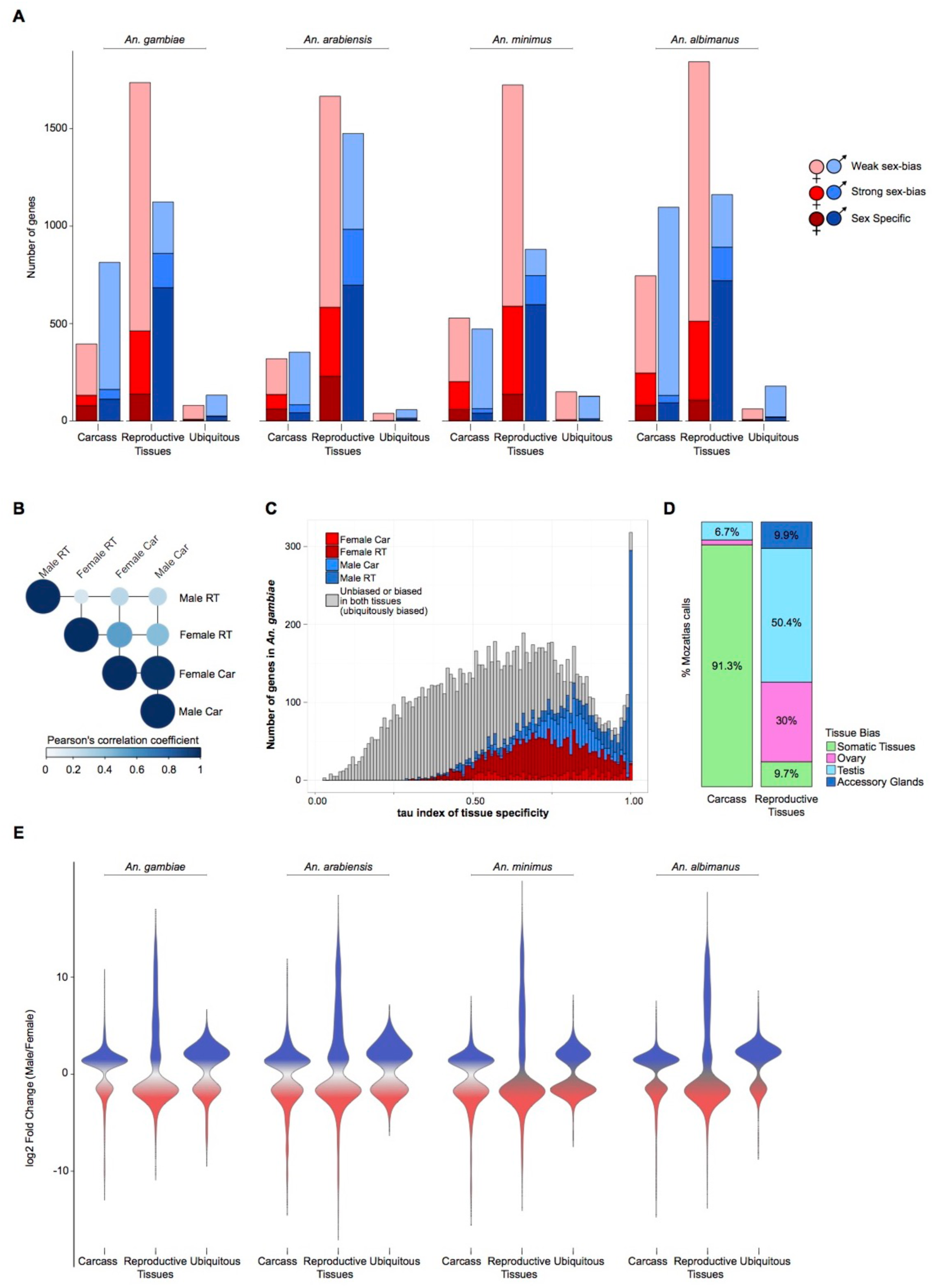
Expression divergence rates of the four-species orthologs. **A)** Median standard deviations of expression ratios (male/female) of carcass and reproductive tissue male-biased (blue) and female-biased (red) amongst the four-species orthologs. **B)** Scatterplots of sex-biased genes of all species in the carcass and the reproductive tissues, showing the relationship of gene expression levels (in log2 FPKM) and expression divergence (standard deviation). **C)** Density distribution plots of gene expression levels (in log2 FPKM) from all four species combined separately for genes with high (SD>1) and low (SD<1) levels of expression divergence. **D)** The number of genes that show sex and/or tissue bias compared to their *An. gambiae* orthologs increases with evolutionary distance. Wilcoxon rank-sum test is used to test for significant differences when comparing sex-bias gene expression categories between the sexes (blue asterisk) or to unbiased genes (gray asterisk) and indicates *P <* 0.001 (**), *P< 0.05 (*).*

The correlation between SD and gene expression levels (FPKM) of sex-biased genes irrespective of their sex-bias category in both tissues was weak but positive (average *R*^2^ = 0.0825, Figure 3B). We noted however, that there is an inverse relationship between median SD and overall gene expression levels (FPKM) among the sex-bias categories, e.g. female-specific genes in the carcass have higher levels of SD but also have lower expression levels than strongly- or weakly female-biased genes (Figure 3A and Sup. Figure3B). To rule out that high expression divergence was not an artifact of low gene expression, we first split the sex-biased genes into subsets of either rapidly and slowly evolving based on a SD value higher or lower than 1, respectively. If high SD rates were predominantly driven by noisy fold change ratios resulting from low gene expression levels we expected that the subset of highly divergent genes (SD>1) (Zhang et al. 2007) would show significantly lower levels of expression. With the exception of the male reproductive tissues, we observed the opposite: that is the expression spectrum of genes with SD>1 shifted to higher expression levels in comparison to the less divergent genes, with significant differences between median levels between the two subsets (Wilcoxon rank-sum test minimum p< 0.05) (Figure 3C). To confirm this, we also divided the entire gene expression range in each tissue and sex into three subsets – low, medium and high expression - and compared the median SD rates of each of the sex-bias categories within these defined ranges of expression (Sup. Figure4). These results confirmed that genes showing stronger sex-bias displayed higher levels of expression divergence, regardless of their levels of gene expression. Qualitatively, reversal of sex-biased expression (e.g. male to female) between species occurs infrequently (e.g. 7 orthologs between *An. gambiae* to *An. arabiensis* show this behavior) and increases with evolutionary distance between species in both the carcass and reproductive tissues (Figure 3D). Moreover, we observed a similar frequency of sex switching and tissue switching between species (Figure 3D).

#### Sequence divergence

To investigate how rates of sequence divergence relate to sex-biased gene expression, we employed both DNA-level and protein-level measures of divergence calculated across all orthologs from the entire *Anopheles* genus (Neafsey et al. 2014). DNA evolutionary constraints were computed for each orthogroup using the number of non-synonymous substitutions per non-synonymous site (dN) and the number of synonymous substitutions per synonymous site (dS) to quantify dN/dS ratios. Protein evolutionary rates were computed to quantify the relative sequence conservation or divergence among orthologs in terms of their normalized amino acid identities (Waterhouse et al. 2011; Kriventseva et al. 2015). Similar to our findings of expression divergence, we found that female-specific genes of the carcass displayed the highest median levels of sequence divergence among all sex-biased genes. (Figure 4A and B), being significantly higher than both male-specific genes and unbiased genes (Wilcoxon rank-sum test p<0.001) in all species. In the reproductive tissues, all three female categories consistently evolved faster than unbiased genes in all species (Wilcoxon rank-sum test p<0.001), and in the males, only the sex-specific genes displayed mean dN/dS ratios and evolutionary rates significantly higher than unbiased genes consistently in all species (Figure 4A and B). To rule out that any of the above filtering steps may have a substantial effect on the observed patterns of sequence divergence, we analyzed DNA-level sequence divergence (dN/dS) of each of the sex-bias categories for each species and found no changes in the overall patterns of divergence between sex-bias categories regardless of the stage of DE analysis or orthology relationships employed (Sup. Figure5). We also tested for an association between expression and sequence divergence among sex-biased genes of all species and found that genes displaying rapid expression divergence (SD>1) also show higher dN/dS ratios compared to those with an SD<1, as has been observed in mammals(Khaitovich et al. 2005) and *Drosophila*(Zhang et al. 2007) (Sup. Figure6).

**Figure 4.**
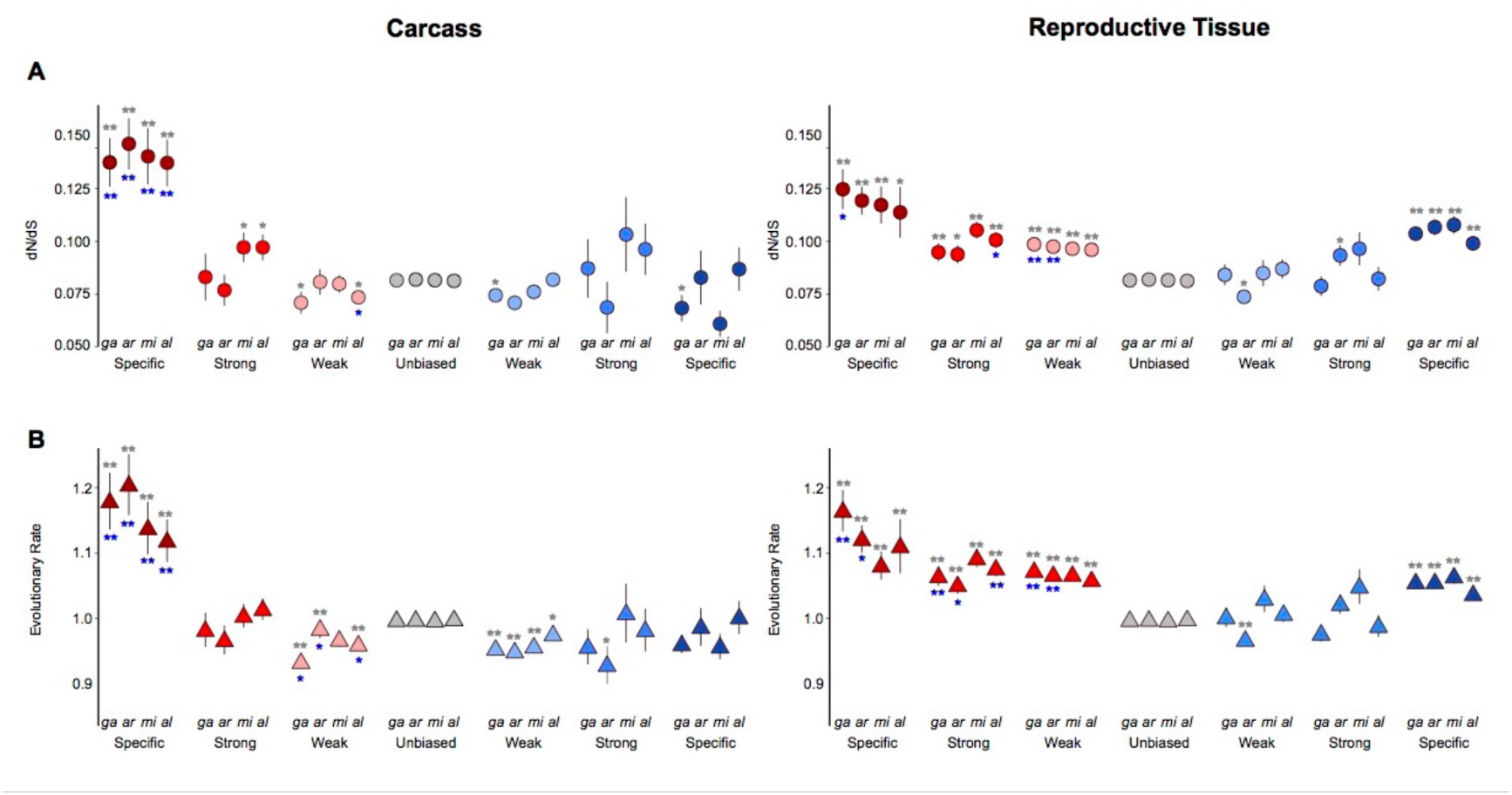
Sequence divergence rates of the four-species orthologs. Average DNA sequence (A) and protein sequence (**B**) divergence levels of the four-species orthologs from each species plotted by the sex-bias expression classification. Wilcoxon rank-sum tests are used to test for significant differences when comparing divergence levels between the sexes (blue asterisk) or between sex-biased categories and unbiased genes (gray asterisk) and indicates *P <* 0.001 (**), *P< 0.05 (*).*

#### Genic turnover

To investigate how patterns of gene gain or loss (turnover) relate to gene expression biases, we assessed the orthology status of all sex-biased genes between the four study species. We first examined the influence of sex-biased expression on the distribution of species-specific genes, as these likely represent cases of high genic turnover either through species-specific gains or widespread losses, or very high levels of sequence divergence that preclude ortholog identification. Average turnover rates were estimated for each species and for each sex-bias expression category by calculating the fraction of species-specific genes from the total number of genes per category (Figure5A). We also quantified the degrees of conservation of each subset of sex-biased genes of *An. gambiae*, ranging from species*-*specific to present in all four species, to gain a more detailed view of gene turnover within the genus (Figure5B). Conservation levels were calculated by diving the number of genes in each conservation and sex-bias class by the total number of genes in the sex-biased category (e.g. number of genes shared in the *gambiae* complex displaying carcass female-specific expression in *An. gambiae* over the total number of carcass female-specific genes). Comparisons of gene turnover rates for each sex-bias expression category to unbiased genes or between the sexes was consistent with our findings on expression and sequence divergence: in both tissues the fraction of species-specific genes increased with the magnitude of sex-biased gene expression (except male-specific in the carcass), and female-biased genes displayed higher or similar levels of turnover to male-biased genes in all species (Figure5A). Furthermore, the extent of conservation throughout the genus in the carcass was lowest for female-specific genes and in reproductive tissues was similar between males and females, with decreasing conservation as the magnitude of sex-bias increases (Figure5B). To test whether this result may have been biased by the inclusion of multigene families, we repeated this analysis removing all orthogroups in which there were any multi-copy orthologs and found a similar result (Sup. Figure7).

**Figure 5.**
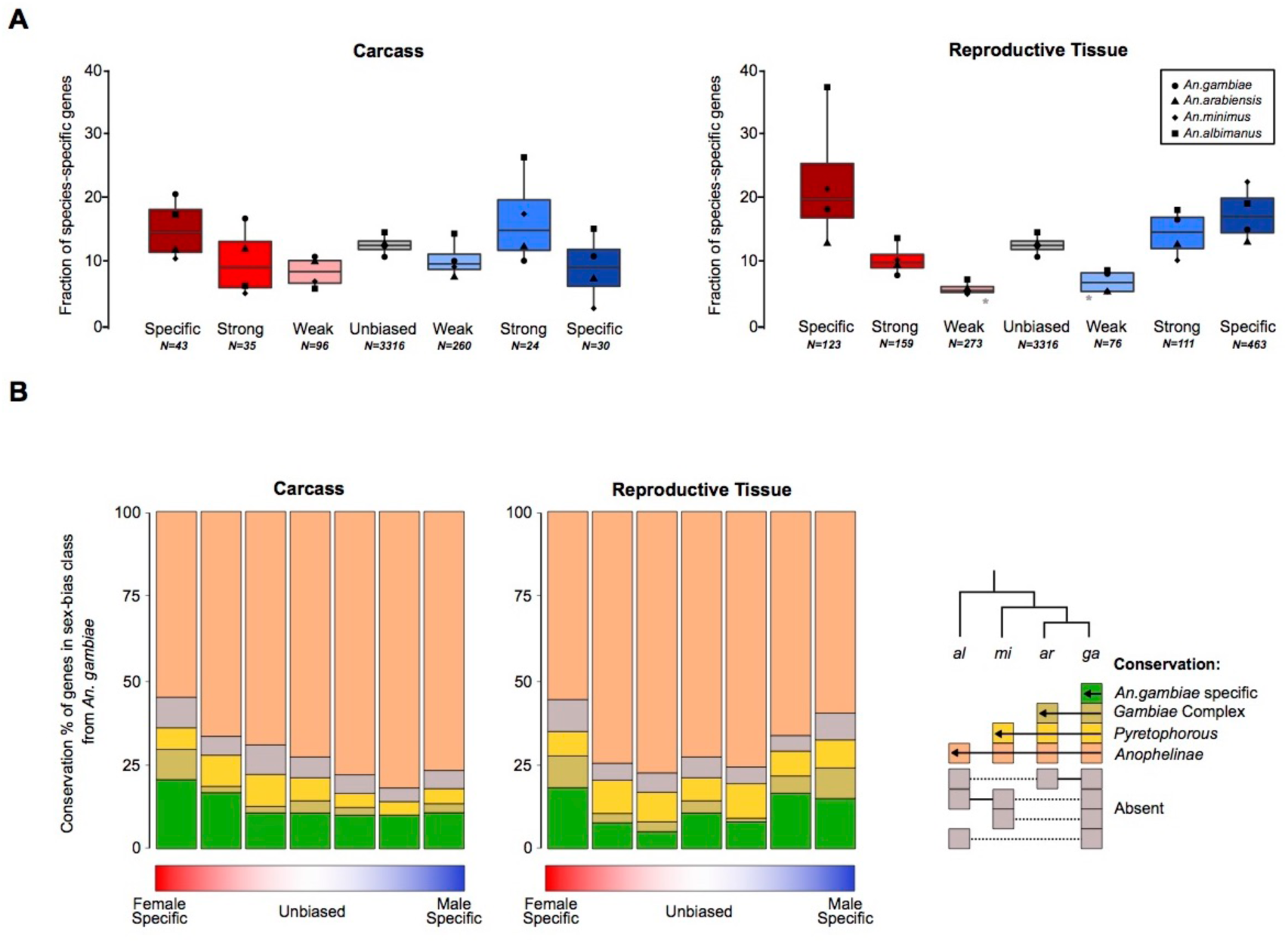
Sex-bias gene turnover rates. (**A**) The fraction of species-specific (orphan) genes is plotted for each sex-bias classification for the carcass and reproductive tissues, either for each species individually (points) or all species (boxplot). Wilcoxon rank-sum test of significance comparing sex-bias gene expression to unbiased genes indicate *p* < 0.05. (**B**) Conservation of *An. gambiae* sex-biased genes across the phylogeny.

### Rapidly evolving sex-biased genes are confined to a few biological functions

To understand the forces that drive the rapid evolution of female-biased genes we investigated the putative functions of the most rapidly evolving and most sex-biased genes in the carcass tissues of *An. gambiae*. We chose this species because of its more comprehensive functional ontologies and the availability of sets of manual community annotations (Neafsey et al. 2014). We found that a limited subset of functional categories describes the functions of nearly two-thirds of sex-biased genes in the female carcass (Figure 6). Function or expression in female salivary glands was the most over-represented annotation with 34% of rapidly evolving female genes falling in this group (Fisher’s exact test for overrepresentation p-value<0.001) (Figure 6A). A large fraction of these genes are known to facilitate blood feeding, e.g. apyrase (AGAP011971), gSG6 (AGAP000150), gSG7 (AGAP008216) and the D7 family of genes (AGAP008278, AGAP008279), because they interfere with host homeostasis, inflammation and immunity (Champagne et al. 1995; Calvo et al. 2006; Isawa et al. 2007; Arcà et al. 2014). Their rapidly diverging sequences and previous work showing that some of these are under strong positive selection (Arcà et al. 2014) lends support to the notion that host immune pressures are a dominant driving force acting on salivary gland genes essential for blood feeding. Salivary glands are also the primary mosquito tissues to become infected with malarial parasites. Therefore genes such as *saglin* (SG1f:AGAP000610) that acts as a binding partner for parasitic TRAP(Ghosh et al. 2009; Upton et al. 2015) proteins, but also 5 more members of this SG1 family (SG1:AGAP000612, SG1c:AGAP000607, SG1e:AGAP000609, SG1b:AGAP000548) whose functions are not yet known, were also among salivary gland genes that are biased towards females and evolving rapidly (Figure 6B). Future studies will be needed to determine whether other SG1-like proteins interact with parasitic counterparts and whether this interaction drives sequence diversification of this family of genes in malaria vectors.

**Figure 6.**
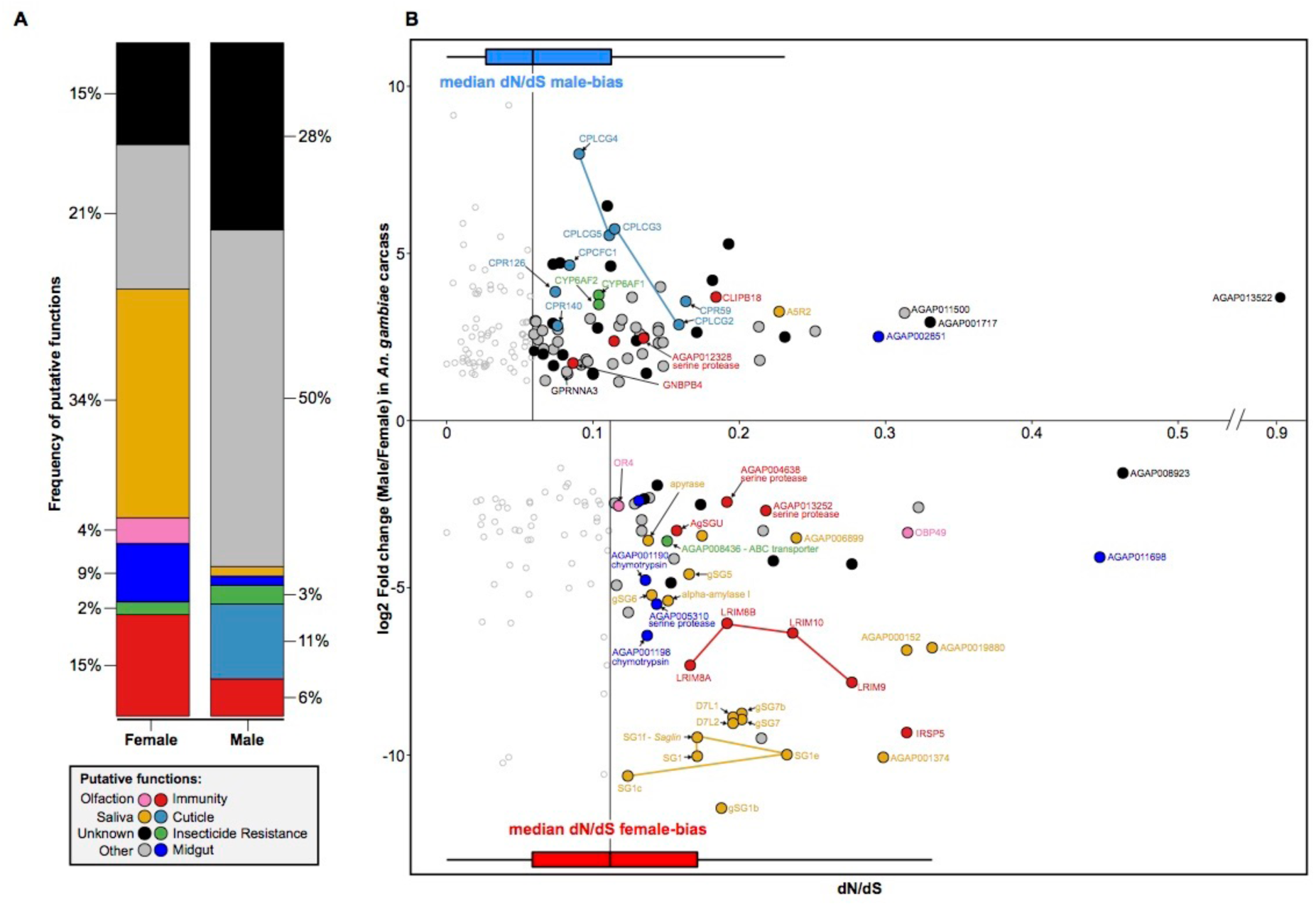
Functional analysis of rapidly evolving and highly sex-biased genes from the carcass of *An. gambiae.* (**A**) Relative proportions of putative functions amongst rapidly evolving sex-biased genes (sex-specific and strongly-biased) in the carcass of *An. gambiae*. (B) Sex-biased expression ratios (in log2 fold change male/female) over sequence divergence rates (in dN/dS) for genes displaying strong-bias or sex-specific expression in the carcass of *An. gambiae*. Horizontal box-plots indicate dN/dS values for female- (red) and male-biased (blue) genes, highlighting the median values of each sex, which were used as cutoff for downstream analysis. Genes with dN/dS value greater than the median of each sex are highlighted with putative functions using GO, orthology-based predicitions and community annotations (colored circles). Empty circles indicate genes evolving below our threshold of sequence divergence.

The second most common functional category was mosquito immunity (15% of female genes; Fisher’s exact test p-value < 0.001) (Figure 6A), and the majority of genes within it have been shown previously to either respond to or influence infection levels (Figure 6B). Among these are cluster on chromosome 2L containing an array of leucine-rich repeat immune proteins (LRIM 8A:AGAP007454, 8B: AGAP007456, 9:AGAP007453 and 10:AGAP007455 (Waterhouse et al. 2010)), of which LRIM9 knockdown increases parasite load 3-fold (Upton et al. 2015; Vlachou et al. 2005). AGAP000151 (IRSP5), whose silencing increases both bacterial and parasite numbers(Dong et al. 2006) and AGAP000570 (AgSGU), which is one of the most abundant peritrophic matrix proteins (Dinglasan et al. 2009) and has important roles in ookinete attachment of *Plasmodium falciparum* (Mathias et al. 2014) were also identified. Of the remaining genes, 4 are expressed in the female midgut and are likely involved in blood meal digestion, 2 are involved in chemosensation (OR4:AGAP011467 and OBP49:AGAP006075) and an ABC transporter (AGAP008436) that is involved in insecticide resistance against permethrin. The putative functions of all other rapidly evolving genes biased towards females are currently unknown or did not fall within these community annotated functional categories.

The number of rapidly evolving male-biased genes of the carcass without predicted functions was higher compared to females (28%) (Figure 6A). Functions related to the insect cuticle, were overrepresented (11% of male genes; Fisher’s exact test p-value < 0.001) including the entire cluster of CPLCG genes on chromosome 3R (CPLCG2: AGAP008445, CPLCG3: AGAP008446, CPLCG4: AGAP008447, CPLCG5: AGAP008449), of which CPLCG3 and CPLCG4 mediate through yet unknown mechanisms insecticide resistance in distinct strains and species of *Anopheles* (Vontas et al. 2007; Awolola et al. 2009). Three additional cuticular proteins, namely CPCFC1 (AGAP007980) and CPR59 (AGAP006829) and CPR140 (AGAP006868) also showed a bias towards males (Figure 6B). No previous report of male biased expression for these genes, or any other cuticle protein, exist. We therefore inspected expression profiles of these genes in published expression data and found that all are expressed females, but are downregulated upon blood feeding, explaining the observed male bias in our data (data not shown). Of the remaining genes, 4 included candidate immunity genes such as GNBPB4 (AGAP002796) and CLIPB18 (AGAP009215), and two cytochrome P450s (CYP6AF1:AGAP011028 and CYP6AF2:AGAP011029), but this category was not functionally enriched among the rapidly evolving highly male biased genes.

### Sex-specific splicing

An exclusive focus on differential expression of genes between males and females may miss a significant portion of sex-biased gene usage, as many processes - as is known for the sex-determination cascade itself, could be regulated at the level of transcript usage or splicing. We therefore expanded the catalogue of sex-biased genes in the four species to include those with sex-specific isoform usage and characterized the degree of conservation of splicing patterns between species. Because the annotation of alternative transcript isoforms in these species is incomplete we first established an isoform-specific gene expression dataset, where we predicted novel isoforms of all known genes (Sup. Figure8). We focused our analysis on orthologs that possessed at least two transcripts that were differentially used between the sexes or tissues, so that each isoform was expressed at least at a 2-fold higher level in the condition in which it was predominant. This stringent cut-off in both directions, together with the requirement for conservation of splicing patterns between species allowed us to identify a credible set of candidate loci that are sex- and/or tissue-specifically spliced in at least two *Anopheles* species. While sex-specific splicing does occur in the carcass, when considering only genes that show high conservation in usage across the genus, the list is restricted to a few genes including the well-known *doublesex* master-switch of the sex determination pathway. We found conserved sex-specific isoform usage between the reproductive tissues of males and females to be at least as common, if not more so than tissue-specific splicing between the carcass and the reproductive tract in either males or females (Figure 7A). In *An. gambiae*, genes that displayed sex-specific isoform usage in both tissues were enriched for functional annotations including behavior, oocyte and embryo development and the regulation of cellular and organismal anatomy (Fisher’s Exact test p-value < 00.01) (Figure 7B). Exemplary transcript structures for genes falling into both classes are shown in Figure 7C and the complete list of genes with predicted alternative transcript usage is listed in Table S3. This analysis confirms that a considerable amount of sex-specific gene usage in the mosquito remains entirely unexplored and we provide candidate loci for future functional analysis.

**Figure 7.**
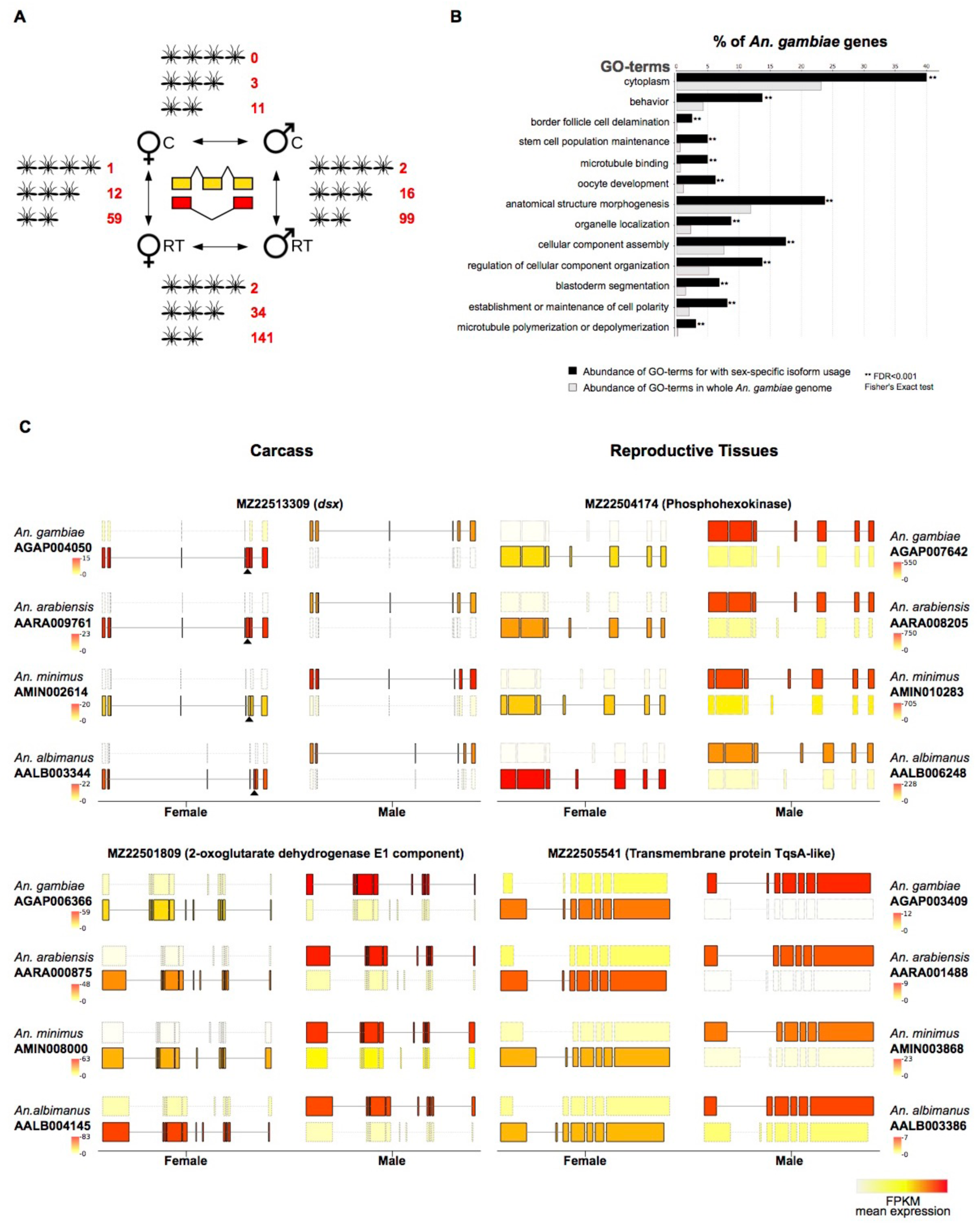
Sex- and tissue-specific differential transcript isoform usage. (**A**) Overview of the number of orthologs displaying conserved alternative isoform usage conserved in 4, 3 or 2 species respectively. (**B**) Gene ontology functional enrichment analysis showing enriched GO-terms in *An. gambiae* among genes displaying sex-specific isoform usage, compared to the entire genome. (**C**) Two examples of sex-specific isoform usage between males and females are shown for carcass (left) and reproductive tissues (right). In each case only the predominant isoforms in the two sexes are shown, with the color of the transcript display transcript expression levels (in FPKM). The female-specific exon of the well characterized *doublesex* (*dsx*) locus is marked with a black triangle. Although visual inspection of splicing pattern highlights consistent use of alternative sex-specific splicing in all 4 species, the *dsx* locus passed the cutoffs only in 3 species. A similar pattern was also observed for a number of other genes and suggests that this analysis is stringent.

### Chromosomal distribution of sex-biased genes

Sex-biased genes have been shown in a number of species to be non-randomly distributed across chromosomes, and typically, in species where males are heterogametic (XY), male-biased genes are underrepresented on the X chromosome (Parisi et al. 2003; Wu and Xu 2003; Sturgill et al. 2007; Ellegren and Parsch 2007). To assess the chromosomal distribution of sex-biased genes in the four *Anopheles* species we calculated the ratio of the number of observed over expected sex-biased genes for each chromosome type (X or autosome) for both tissues. We observed a significant paucity for male RT-biased genes on the X-chromosome in all species (*χ*^2^ test *p*<0.001), but found no similar deficit among male-biased gene of the carcass (Figure 8A). These results extend our previous findings in *An. gambiae* (Magnusson et al. 2012) in two ways: First, they demonstrate that the underrepresentation of male-biased genes on X chromosomes is a conserved feature in the genus. Second, they show that the underrepresentation detected previously using whole-body data is derived from differences in the abundance of sex-biased genes in the reproductive tissues. While the X-chromosomes did show a higher than expected number of female-biased genes in some cases (Figure 8A), this apparent “feminization” was not as significant and not consistently found in all four species (*χ*^2^ test *p*<0.05).

**Figure 8.**
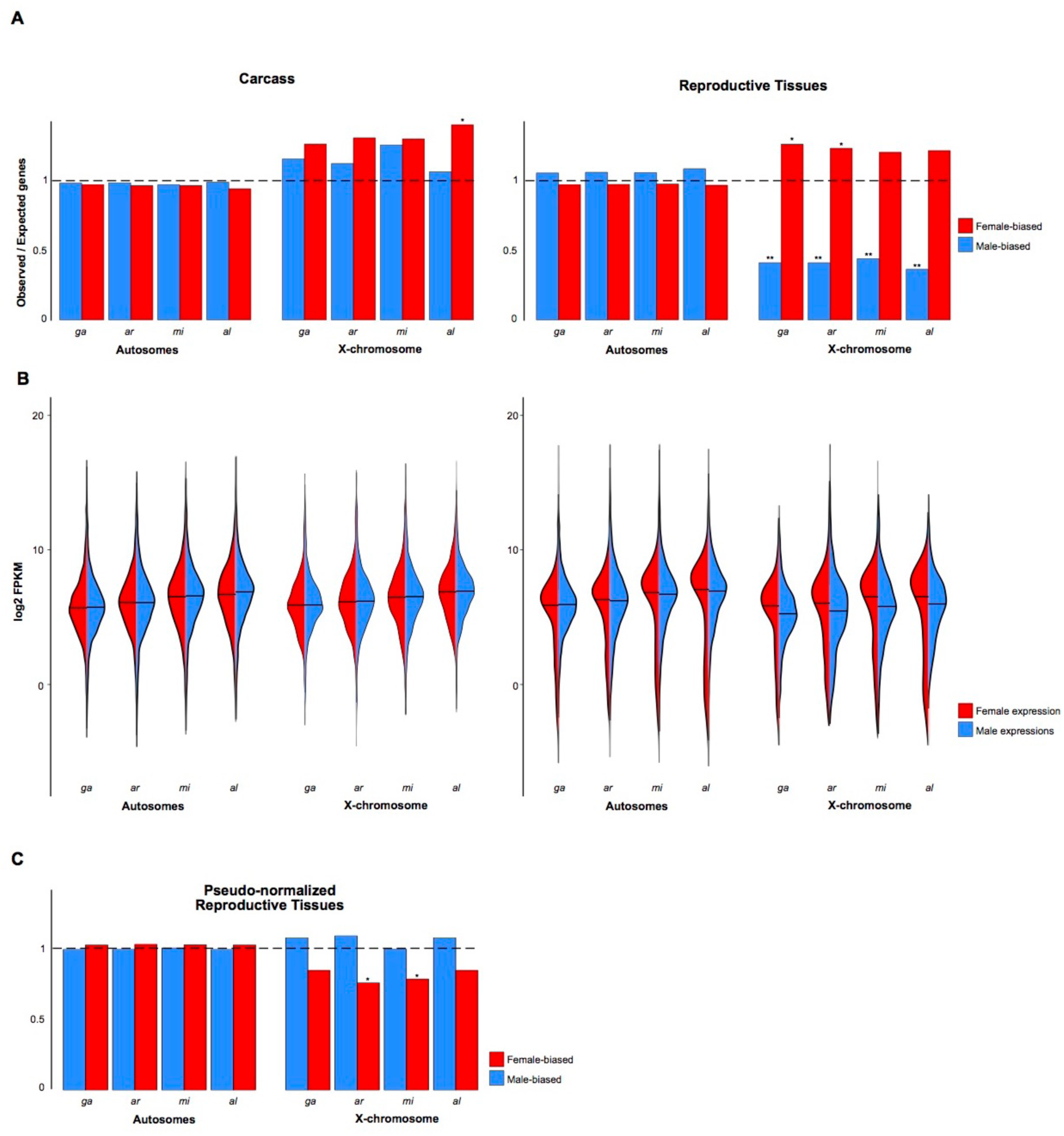
Chromosomal distribution of sex-biased genes. **(A)** Proportions of observed/expected numbers of male-biased (blue) and female-biased (red) genes in the carcass and reproductive tissues located on autosomes or the X chromosomes of each species. The horizontal dashed line indicates an equal number of observed and expected genes and ratios greater or lower than this indicate an enrichment or depletion, respectively. Asterisks indicate *P*<0.05 (*) and *P*<0.001 (**) of a *χ*^2^ test comparing ratios between the X chromosome and the autosomes. **(B)** Violin plots of expression levels (in log2FPKM) of all genes in the female (red) or male (blue) carcass and reproductive tissues for autosomes and X chromosomes. (**C**) Pseudo-normalization of X-chromosome male expression in the reproductive tissues using the A:X ratio eliminates the demasculinization effect.

In *Drosophila*, the deficit of male-biased genes on the X chromosome has been attributed to chromosome-wide transcriptional regulation(s) in the male germline, namely lack of dosage compensation and transcriptional suppression of the X - in a process similar to meiotic sex chromosome inactivation (MCSI), except that the suppression appears to be initiated in pre-meiotic cells (Meiklejohn et al. 2011; Meiklejohn and Presgraves 2012). In combination, these processes result in a significant reduction in gene expression along the entire X chromosome in the male germline, observed as a paucity of male-biased genes on the X chromosome, when expression is compared to females or other tissues. It has been argued that the use of the term “evolutionary X demasculinization” may thus be misleading, because it implies that selective forces, such as sexual antagonism, drive genes beneficial for males off the X (Meiklejohn et al. 2011; Meiklejohn and Presgraves 2012). This argument is supported by the finding that testis-specific genes are in fact not underrepresented on the X of *Drosophila* (Meiklejohn and Presgraves 2012). Instead, testis-specific genes have recruited strong cis-regulatory elements to overcome the transcriptional silencing (Landeen et al. 2016). In *An. gambiae,* we have previously shown that transgenic reporters are transcriptionally suppressed specifically in the mosquito testis (~10-fold) when inserted on the X chromosomes (Magnusson et al. 2012). However, we did not see any evidence for chromosome-wide transcriptional regulation in that study - that is genes on the X chromosome were expressed at similar levels overall between males and females, or between X chromosomes and autosomes in males (Magnusson et al. 2012).

To determine whether there is X chromosome transcriptional regulation in the male reproductive tissues in *Anopheles*, we evaluated chromosome-wide expression levels between males and females - reasoning that the failure to detect this in our previous work may have resulted from the lack of tissue-level expression. Indeed, we found that the median expression of all genes on the X chromosomes is on average 0.3 fold lower in males compared to females in the reproductive tissues in all four species (Figure 8B). In comparison, median expression of X genes in the carcass tissues, and autosomal genes in both tissues were not significantly different between the two sexes (Figure 8B). To confirm this, we calculated the autosome to X chromosome (A:X) ratio of median gene expression in the male reproductive tissues, and as a control for the carcass. We found that in all species the A:X ratio ranged between 1.61 and 1.92 in the male reproductive tissues (Table 1).

**Table 1:** Gene counts, expression data and autosome:X chromosome ratio in the four species for carcass and reproductive tissues. Indicated *p* values were calculated using the Wilcoxon rank sum test.

|  | Number of genes |  | Female |  |  |  | Male |  |  |  |
| --- | --- | --- | --- | --- | --- | --- | --- | --- | --- | --- |
|  | X | Autosomes | X median FPKM | Autosome median FPKM | AA:XX ratio | *P* value | X median FPKM | Autosome median FPKM | AA:X ratio | *P* value |
| **Carcass** |  |  |  |  |  |  |  |  |  |  |
| *An. gambiae* | 1097 | 11923 | 53.7 | 45.7 | 0.85 | 0.0021 | 55.9 | 48.6 | 0.87 | 0.003 |
| *An. arabiensis* | 1278 | 12399 | 49.6 | 41.9 | 0.84 | 0.0003 | 50.1 | 42.7 | 0.85 | 0.001 |
| *An. minimus* | 1217 | 11561 | 59.9 | 58.5 | 0.97 | 0.3091 | 66.5 | 66.6 | 1.00 | 0.523 |
| *An. albimanus* | 1024 | 7776 | 90.1 | 69.0 | 0.76 | 3.415e-07 | 97.2 | 83.3 | 0.86 | 0.002 |
| **Reproductive Tissues** |  |  |  |  |  |  |  |  |  |  |
| *An. gambiae* | 1097 | 11923 | 49.6 | 50.3 | 1.01 | 0.581 | 34.8 | 56.2 | **1.61** | < 2.2e-16 |
| *An. arabiensis* | 1278 | 12399 | 29.0 | 31.3 | 1.07 | 0.275 | 23.6 | 45.4 | **1.92** | 3.89e-14 |
| *An. minimus* | 1217 | 11561 | 43.9 | 49.7 | 1.13 | 0.852 | 38.6 | 72.6 | **1.88** | < 2.2e-16 |
| *An. albimanus* | 1024 | 7776 | 41.9 | 34.5 | 0.82 | 0.178 | 49.5 | 85.9 | **1.73** | 3.89e-14 |

Together, these results suggest that in the male reproductive tissues of *Anopheles,* X-chromosomes undergo chromosome-wide transcriptional regulation, similarly to *Drosophila* (Landeen et al. 2016). We tested whether such X-specific regulation can explain the underrepresentation of male-biased genes on the X chromosomes by simulating the absence of X-regulation in the male reproductive tissues. We first normalized the raw read counts of male X chromosome genes with the respective A:X expression ratio that we computed for each species. We then repeated the DE analysis to re-classify sex-biased gene expression. Using this model that corrects for X transcriptional regulation, the ratios of observed/expected genes no longer support the underrepresentation of male-biased genes on the X chromosomes in the male reproductive tissues of these species (Figure 8C). Finally, we also assessed the chromosomal distribution of genes specifically expressed in the male reproductive tissues. To do this, we calculated the index of tissue specificity (*τ*) for each gene and selected genes with a *τ* index > 0.95. When we tested for evidence of X demasculinization for RT-specific genes using the restricted range of *τ* >0.95, we found no significant paucity on the X compared to autosomes (*χ*^2^ test *P*=0.26). This is consistent with our results indicating that the paucity of male-biased genes on the X chromosome in reproductive tissues results from X-chromosome transcriptional regulation, its effect on DE analyses, and is not likely a result of selective forces such as sexual antagonism.

## Discussion

Studies in many different organisms investigating the relationship between expression patterns and rates of molecular evolution have consistently found evidence for complex selective pressures acting differentially on male- and female-biased genes. Typically, genes biased towards males evolve more rapidly than female biased genes, with those expressed during spermatogenesis showing the fastest rates of evolution in the genome. This accelerated evolution can occur at the sequence level, the expression level and the gene turnover level. Amazingly, this pattern is upheld from primates (Khaitovich et al. 2005) to worms (Cutter and Ward 2005), i.e. even in species in which the two sexes coexist within a single individual.

Our findings suggest that *Anopheles* mosquitoes do not follow this common pattern. In all four sequenced species, genes expressed specifically in female carcass tissues consistently display the highest levels of DNA- and protein sequence divergence, expression divergence, and genic turnover. By exploring the putative functions of these genes using manual annotations and expression data, we propose that this pattern is driven by the strong selective forces acting on female-specific phenotypic characters that facilitate blood-feeding and thus underlie vectorial capacity. Nearly two thirds of female genes that are rapidly evolving in *An. gambiae* have predicted functions involving host seeking, blood feeding, digestion, and mounting an immune response to control pathogens. Genes with functions in the female salivary glands, whose products are either injected into the host or are located in the interface with the ingested blood meal within salivary gland tissues, are significantly overrepresented among rapidly evolving female genes (Figure 6). This finding reinforces the notion that both host pressures and mosquito-pathogen interactions are principal forces acting on the evolution of mosquito genomes (Neafsey et al. 2014; Arcà et al. 2014). Our data also highlight that male biology is not under similar selection pressures as those of females but the less-well studied biology of the male somatic tissues makes it harder to establish a link between rapidly evolving male genes in the carcass and their role in male-specific processes or behaviors - for example their aggregation and mating in swarms.

Intriguingly, we found that genes biased towards male reproductive tissues did not display faster rates of evolution than their female counterparts, as has been observed so consistently in other species. Generally, increased evolutionary rates: among male biased reproductive genes are explained by selection arising from male-male competition (Zhang and Parsch 2005; Pröschel et al. 2006; Butler et al. 2007) and/or reduced pleiotropy of genes expressed in male reproductive tissues (Zhang et al. 2007; Meiklejohn and Presgraves 2012; Tripet et al. 2003; Pondeville et al. 2008). Our analysis on tissue specificity in *Anopheles* shows that male RT-biased genes display the highest overall tissue specificity of all sex- and tissue bias categories (Figure 2C and Sup. Fig2), so increased pleiotropy of male RT genes in these mosquitoes is unlikely to explain this result. More likely, the fact that *Anopheles* females typically mate once during their lifetime under natural conditions (Tripet et al. 2003) results in the absence of selection on sperm competition that typically drives increased rates of evolution in male RT genes in polygamous species. Males of several *Anopheles* species including *An. gambiae* and *An. arabiensis* produce a mating plug that is deposited in the female reproductive tract during mating, enhancing sperm storage (Rogers et al. 2009) and modifying female physiology (Pondeville et al. 2008; Baldini et al. 2013) such that females are no longer receptive to further matings. It will be interesting to compare rates of evolution of male and female biased genes in other mosquito species where females are polyandrous (Boyer et al. 2012) or other insect taxa where both sexes bloodfeed (Yuval 2006).

In addition to sex-biased gene expression, differential isoform usage between sexes, resulting from alternative splicing or alternative transcription start sites, can also account for sexual dimorphism. This area remains completely understudied due to the difficulty of extracting isoform-level data from whole genome expression data. In mosquitoes only two genes, *doublesex* and *fruitless*, initially characterized in *Drosophila*, are known to behave in this manner (Scali et al. 2005), but no systematic analysis has been performed to date. Making use of novel tools for isoform-level differential expression analysis and combining it within a comparative analysis across four malaria mosquito species, we have catalogued conserved genes that display sex-specific isoform usage in the carcass and the reproductive tissues. These candidate genes form the basis for future functional investigations into processes such as sex-determination or behavior, for example by the depletion of sex-specific transcripts or exons using CRISPR/Cas9.

Finally, we have also used our expression data to re-examine evidence of the non-random chromosomal distribution of sex-biased genes in *Anopheles* mosquitoes and to begin to characterize its underlying causes. We have previously shown using whole-body expression data from adults and larvae that the X chromosome of *An. gambiae* has a significant paucity of male-biased genes, and that transgenes that are normally expressed strongly in the male testis are transcriptionally suppressed in sperm when the transgenic construct is located on the X chromosome, in a process resembling MSCI (Magnusson et al. 2012). The results from the present study are consistent with our previous work, and reveal that the paucity of male-biased genes on the X chromosome is a conserved feature in the genus. We show that this underrepresentation is specific to reproductive-tissues in all species and that in the carcass there is no significant difference in the distribution of sex-biased genes between chromosomes (Figure 8A). Through exploring the potential forces driving this deficit of male-biased genes, we find that X chromosomes must differ significantly in their chromosome-wide transcriptional regulation between tissues: in the carcass, the difference in ploidy of the X between males and females, or the X and autosomes in males, does not translate into chromosome-wide gene expression differences, suggesting that these tissues are actively dosage compensated - probably through X chromosome hypertranscription in males as in *Drosophila*. However, in the male reproductive tissues, we find that gene expression from the X chromosome is on average 0.3 fold lower in males compared to females and on average 1.8-fold lower compared to gene expression from the autosomes for all four species (Figure 8B and Table 1). This finding is congruent with the results from the Baker and Russell study, which focusing on *An. gambiae* with tissue atlas data showed that X-linked genes are expressed at half of autosomal levels in the male testis (Baker and Russell 2011). We propose however, that the transcriptional regulation occurring in the male reproductive tissues on the X chromosome is directly responsible for the deficit of male-biased genes in the male reproductive tissues, and is not merely a contributing factor favoring X-to-autosome retro-movement of male genes leading to the evolutionarily demasculinization of the X. We show this by computationally simulating the absence of the observed X chromosome transcriptional regulation, which results in effectively eliminating the underrepresentation of male-biased genes on the X. Furthermore, when analyzing the chromosomal distribution of tissue specific genes, which are not similarly sensitive to DE-biases that assume equal ploidy or chromosomal transcriptional regulation between conditions, we find no support that male reproductive tissue specific-genes being underrepresented on the X chromosome, again similarly to *Drosophila* (Meiklejohn et al. 2011). Taken together, these results indicate that the X chromosomes of *Anopheles* mosquitoes are in effect “transcriptionally demasculinized” in the male germline by chromosome-wide transcriptional regulation that leads to reduction of X chromosome gene expression. In the absence of mutant strains or markers for the chromatin-modifications that govern the levels of X transcription in the male germline, it is currently difficult to quantify the individual contributions of the lack of X-dosage compensation or X-suppression (via processes like MSCI), especially since these have similar outcomes. The ~1.8-fold reduction in gene expression between X chromosomes and autosomes in the male reproductive tissues in our data is similar to levels observed in testis of *Drosophila* (Meiklejohn et al. 2011; Meiklejohn and Presgraves 2012), where genetic and cytological data support both the presence of transcriptional suppression on the X chromosome and the absence of dosage compensation. We anticipate that the presence of non-germline tissues within our male reproductive tissue samples - the accessory glands and the somatic sheath enveloping the testes, likely diminish the fold difference in expression of the X chromosome, as we would expect a nearly 10-fold difference to arise from X chromosome suppression alone (Magnusson et al. 2012). Dissecting the mechanisms and functions of this transcriptional regulation will likely reveal important insights into the evolution of sex-chromosomes in *Anopheles,* including addressing whether similar transcriptional regulation occurs on the Y chromosomes, whose content and evolutionary history we have recently described (Hall et al. 2016).

## Methods

### RNA isolation and RNA sequencing

Total mRNA was extracted from sexed samples derived from MR4/BEI colonies of the four mosquito species, namely *An. gambiae* G3, *An. arabiensis* Dongola*, An.minimus MINIMUS1* and *An. albimanus* STECLA. Samples comprised 3-4 day old adult males and 48 hour blood-fed females from a mixed cage. Individuals were dissected to obtain the reproductive tissues, which consisted of ovaries and lower reproductive tract for females and testis and accessory glands for males (Figure 1C). The remaining body including the head, thorax and abdomen were used as the carcass samples. Three replicates were collected for each sample. mRNA was prepared for sequencing using the Illumina mRNA-Seq Sample Preparation kit and samples were paired-end sequenced using 100bp reads. The resulting data for *An. arabiensis*, *An. minimus* and *An. albimanus* have been submitted to the NCBI SRP083856. *An. gambiae* data were previously published to the NCBI SRP045243.

### Differential expression analysis

TopHat2 (Kim et al. 2013) was used to align paired-end RNA-seq data from each species on the reference genomes (version 4.2 for *An. gambiae*, 1.3 for *An. arabiensis*, 1.3 for *An. minimus* and 1.3 for *An. albimanus*), which were retrieved from Vectorbase (Giraldo-Calderón et al. 2015). Mapping was performed using default settings and the number of mapping reads for each gene were extracted using HTSeq (Anders et al. 2014). Sex-biased differential expression analysis was performed using the DESeq-2 (Anders and Huber 2010). Two consecutive DE analyses were conducted: first testing for “male-female” gene expression-bias between the sexes of the same tissue, and then a “carcass-reproductive” bias between tissues of the same sex, at a FDR<0.05 between conditions (Sup Figure 1). Sex-biased genes that showed significant and concordant tissue biases were classified into categories reflecting both traits. When genes were significantly sex-biased in both tissues towards one sex, but there was no significant bias between tissues of that sex, these were classified as ubiquitously sex-biased. Genes displaying conflicting sex- and tissue-bias (e.g. male-bias between RT samples but Car-biased within males) were not included in the final sex-bias categories. To categorize genes according to the magnitude of sex-biased expression, sex-biased genes were divided into three mutually exclusive categories. (1) Sex-specific: when a gene was detected with more than 20FPKM in one sex but less than 10FPKM in the other, (2) strong sex-bias: with a sex expression ratio higher than 5 and (3) weak sex-bias: with a sex expression ratio higher than 2 but lower than 5. The tissue specificity index, τ, was calculated for each gene using the formula

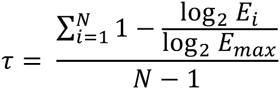

where *N* is the number of tissues, *E*_*i*_ is expression in tissue *i,* and *E*_*max*_ is the maximal expression level for that gene across all samples (Yanai et al. 2005; Larracuente et al. 2008; Meiklejohn and Presgraves 2012). τ ranges from 0 to 1, with higher τ values indicating greater tissue specificity.

### Evolution of sex-biased genes

We retrieved OrthoDB-delineated (Kriventseva et al. 2015) orthology relationships of all genes between the four species (Neafsey et al. 2014) and calculated expression, sequence and gene turnover rates of sex-biased and unbiased genes of all species. Expression divergence and sequence divergence levels were calculated focusing on the 7400 four-species orthologs. Expression divergence was estimated by computing the median standard deviation (SD) of the log2 FC between the four orthologs, for each sex-bias category (including unbiased), separately for carcass and reproductive tissues. Sequence divergence levels were calculated both at the DNA level (dN/dS) and the protein level (evolutionary rate) for each species and each sex-bias category. We used the Wilcoxon test to compare divergence estimates between unbiased and sex-bias categories, or between identical sex-biased categories in the two sexes. The frequency of species-specific genes was used to infer gene turnover rates for each sex-bias category, species and tissue. Furthermore, genic turnover rates were also calculated by dividing the number of genes in each sex-bias category displaying distinct conservation levels in the genus (e.g. species-specific or present in all species), by the total number genes in that category.

### Functional analysis

Gene Ontology (GO) (Ashburner et al. 2000; Blake et al. 2015) and InterPro (Mitchell et al. 2015) terms were retrieved from Vectorbase (Giraldo-Calderón et al. 2015) for each species. We expanded these ontology datasets using Blast2GO (Conesa et al. 2005) using the non-redundant NCBI protein database. For the functional analysis of *An. gambiae* sex-biased genes we also incorporated manual functional annotations provided generously from the community that included chemosensation (gustatory and olfactory), neuropeptide hormones, immunity (recognition; signal transduction; cascade modulation; effectors), insecticide resistance, epigenetic modification, cuticle formation, as well gene expression datasets in the female salivary gland and male accessory gland.

### Chromosomal distribution and expression of sex-biased genes

Gene-to-chromosome assignments are currently only available for *An. gambiae.* To identify the chromosomal origin of scaffolds in the other species, scaffolds were assigned to a chromosomal arm if more than half of the genes on a scaffold had orthologs in *An. gambiae,* and of these, if two thirds had hits to the same chromosomal arm of *An. gambiae*. To assess the reliability of this approach we mapped whole genome sequencing (WGS) data based on Illumina pair end reads from male and female pools of *An. arabiensis* (NCBI SRA project SRP044010) to the *An. arabiensis* scaffolds and calculated the ratio of mapping reads from females over males for each scaffold. X-linked scaffolds were distinguished from autosomal scaffolds when scaffolds displayed a mapping ratio higher than 1.48, which is the mean ratio of all scaffolds plus one SD of the ratio (Sup. Figure8). We found that all 1200 orthology predicted X-linked genes in *An. arabiensis* were derived from scaffolds with female-biased coverage in mapping DNA reads. For simplicity, autosomal arms were combined for all subsequent analysis, as re-arrangements of autosomal arms within the genus have been documented (Neafsey et al. 2014). We calculated the expected number of male/female-biased genes for each chromosome by multiplying the total number of male/female-biased genes in the genome by the proportion of all expressed genes per chromosome. A ratio of observed over expected of 1 would imply there is a random distribution of genes, while values higher than 1 indicate an enrichment, and values lower than 1 indicate a paucity. A *χ*^2^ test was performed to determine whether the ratio of observed over expected numbers of male- and female-biased genes were significantly different between the autosomes and the X-chromosome for the carcass and reproductive tissues for each sex. To assess whether X chromosomes were dosage compensated in both tissues we calculated the ratio of autosome/X-chromosome median gene expression. To understand if the apparent demasculinization of the X chromosome was due to a lack of dosage compensation in the male reproductive tissues, we “pseudo-normalized” the numbers of mapping raw read counts for X-linked genes in the male reproductive tissues of each species by multiplying them by the A:X ratio from each species. We then re-run the DE pipeline to re-classify sex-biased expression. Using this modeled dosage compensation, we calculated the ratios of observed/expected genes finding no evidence of an underrepresentation of male-biased genes on the X chromosomes of each species.

### Analysis of sex-specific isoform usage

RNA-seq reads were aligned separately to the aforementioned downloaded genomes using HISAT (Kim et al. 2015). We then used Stringtie (Mihaela Pertea 2015) to predict novel isoforms of known genes separately for each of the 4 species across the samples. For each species predicted isoforms for all genes were merged into a combined geneset using Cuffmerge (Trapnell et al. 2010). Expression of all isoforms was then quantified using Stringtie. We used the Ballgown (Frazee et al. 2015) suite to determine significantly different isoform expression using a cutoff of p<0.05 and a mean expression greater than 1 FPKM across the samples in a way that each gene was required to have at minimum one male and one female biased isoform with a log2 fold-change greater than 1 between the two sexes.

## Data Access

All RNA-seq data can be downloaded from NCBI SRAs under accession numbers SRP083856 for *An. arabiensis*, *An. minimus* and *An. albimanus* and SRP045243 for *An. gambiae*. The DNA WGS data used for assessing X chromosome linkage of scaffolds in *An. arabiensis* can be downloaded from NCBI SRA under accession number SRP044010.

## Acknowledgements

We would like to thank Scott Cornman, Sara Mitchell, Adam Jenkins, Michael Riehle, Craig Wilding, Xiofan Zhou and Jose Ribeiro for providing us with manual community annotations of *An. gambiae* gene functions, and Paul Howell at the MR4/BEI for support on rearing of the mosquito species. This study has received funding from the European Union’s Seventh Framework Programme (FP7 2007-2013 Infravec) under the G.A. no 228421. This study was funded by the European Union’s Seventh Framework Programme (FP7 2007-2013) Marie Curie Actions cofund (Project I-Move) under GA no. 267232. This study was funded in part by a grant from the Foundation for the National Institutes of Health through the Vector-Based Control of Transmission: Discovery Research (VCTR) program of the Grand Challenges in Global Health initiative of the Bill & Melinda Gates Foundation. P.A.P. was supported by a Rita Levi Montalcini award from the Ministry Education, University and Research (MIUR – D.M. no. 79 04.02.2014). This study was funded by the European Research Council under the European Union’s Seventh Framework Programme ERC grant no. 335724 awarded to N.W. R.M.W. was supported by Marie Curie International Outgoing Fellowship PIOF-GA-2011-303312. M.K.N.L. was supported by an MRC Career Development Award (G1100339) and by the Wellcome Trust (098051).

## Author Contributions

F.P., N.W., R.M.W. and P.A.P. performed all analysis and wrote the paper with help from T.N. and M.L. TN and ML contributed to the experimental design of the study. A.C., R.D. and T.P. performed tissue dissections and RNA-seq library preparations.

## Supplementary Figure Legends

**Supplementary Figure 1.** Flow chart and results of the two-step differential expression analysis in the four species: first testing for a sex-bias between tissues (e.g. male Car vs female Car) and secondly a tissue-bias within each sex (e.g. male Car vs male RT) using a false discovery rate<0.05 between conditions. The two DE analysis were combined and only genes displaying concordant classifications in both analyses were classified as sex-biased. Numbers represent the amount of differentially expressed genes for each species in each step.

**Supplementary Figure 2.** Distribution of *τ* values of tissue specificity in *An. arabiensis, An. minimus* and *An. albimanus.* Colors indicated sex- and tissue-biased classification of the DE pipeline. Genes with high *τ* values are enriched for male-biased reproductive tissue genes in all three species.

**Supplementary Figure 3.** Expression divergence levels for each study species and relationship between gene expression levels and sex-bias classifications. (**A)** Levels of expression divergence (in standard deviation SD) of carcass and reproductive tissue male-biased (blue) and female-biased (red) genes, shown for each species individually. Wilcoxon rank-sum test of significance comparing sex-bias gene expression categories between the sexes (blue asterisk) or to unbiased genes (gray asterisk) and indicated *P* < 0.001 (**), *P*< 0.05 (*). **B)** Levels of gene expression (in FPKM) of all four species and their relationship to the sex- and tissue-bias classification.

**Supplementary Figure 4.** Expression divergence levels are not influence by gene expression levels. For each tissue and sex, the spectrum of gene expression of all genes is divided into three subsets (lowly expressed, medium expressed and highly expressed). For each subset, the standard deviation (SD) was calculated for all four species separately for sex-specific, strongly- and weakly-biased genes. *N* indicates the number of genes in each subset. In yellow are shown the FPKM values selected to create the three subsets

**Supplementary Figure 5.** DNA sequence divergence rates (dN/dS) of sex-biased genes with increasing implementations of filtering does not change overall patterns. Tow row: sex-biased genes without tissue bias classification (1^st^ DE analysis), second row: inclusion of results from the 2^nd^ DE analysis, where tissues biases within each sex are evaluated. The lower three panels show dN/dS levels for each sex- and tissue-biased classification but including only certain orthology relationships: many-to-many; one-to-one at least two species and finally the one-to-one four-species orthologs. The observation of higher sequence divergence for female-specific genes of the carcass remains consistent in all stages of filtering.

**Supplementary Figure 6.** Relationship between sequence and expression divergence rates of sex-biased genes in the four species. Mean DNA sequence divergence levels (dN/dS) increase amongst genes that display high levels of expression divergence (SD>1, darker colors), compared to genes showing lower levels of expression divergence (SD<1, lighter colors).

**Supplementary Figure 7.** Conservation levels of *An. gambiae* sex-biased genes across the *Anopheles* phylogeny, allowing only one-to-one relationships among orthologs. As in figure Figure 5B in which all relationships among orthologs are considered, the fraction of species-specific genes increases with the magnitude of sex-biased gene expression (except male-specific in the carcass), and female-biased genes displayed higher or similar levels of turnover to male-biased genes in all species and in both tissues.

**Supplementary Figure 8.** Improving the alternative isoforms for non-*An. gambiae* species is required for alternative isoform usage analysis. Shown is the number of loci that contain from 1 to 10 annotated isoforms in (**A**) the reference gene-sets of *An. gambiae* (GAM), *An. Arabiensis* (ARA), *An. minimus* (MIN) and *An. albimanus* (ALB) as well as in (**B**) the expanded gene-sets of the same species generated from RNA-seq data from this study.

**Supplementary Figure 9.** Reliability of synteny-based predictions to assign chromosomal origin of genomic scaffolds in *An. arabiensis.* Ratio of mapping female and male WG reads of *An. arabiensis* against assembled scaffolds. Horizontal line at 1.48 shows the mean of the mapping ratio of all scaffolds plus one SD, which was used to distinguish X-linked scaffolds. Each point represents a scaffold in *An. arabiensis* and its position on the X axis is based the number of genes it contains. Scaffold colors show results from orthology-predictions and/or ratio-based decisions of chromosomal location (green – autosomal scaffolds; red - X-linked scaffolds based on orthology; purple – X-linked scaffolds based on ratio but insufficient number of genes for orthology predictions (excluded from the analysis), grey – scaffold with no annotated genes precluding orthology predictions to *An. gambiae*).

